# Genomic Evidence for Formate Metabolism by Chloroflexi as the Key to Unlocking Deep Carbon in Lost City Microbial Ecosystems

**DOI:** 10.1101/831230

**Authors:** Julia M. McGonigle, Susan Q. Lang, William J. Brazelton

## Abstract

The Lost City hydrothermal field on the Mid-Atlantic Ridge supports dense microbial life on the lofty calcium carbonate chimney structures. The vent field is fueled by chemical reactions between the ultramafic rock under the chimneys and ambient seawater. These serpentinization reactions provide reducing power (as hydrogen gas) and organic compounds that can serve as microbial food; the most abundant of these are methane and formate. Previous studies have characterized the interior of the chimneys as a single-species biofilm inhabited by the Lost City Methanosarcinales, but also indicated that this methanogen is unable to metabolize formate. The new metagenomic results presented here indicate that carbon cycling in these Lost City chimney biofilms could depend on the metabolism of formate by low-abundance Chloroflexi species. Additionally, we present evidence that metabolically diverse, formate-utilizing *Sulfurovum* species are living in the transition zone between the interior and exterior of the chimneys.

**IMPORTANCE:** Primitive forms of life may have originated around hydrothermal vents at the bottom of the ancient ocean. The Lost City hydrothermal vent field, fueled by just rock and water, provides an analog for not only primitive ecosystems but also extraterrestrial ecosystems that might support life. The microscopic life covering towering chimney structures at the Lost City has been well characterized, yet little is known about the carbon cycling in this ecosystem. These results provide a better understanding of how carbon from the deep subsurface can fuel rich microbial ecosystems on the seafloor.

## INTRODUCTION

The towering carbonate chimneys of the Lost City hydrothermal vent field protrude from the Atlantis Massif, a dome of ultramafic rock uplifted from the mantle. These chimneys differ from other deep-sea hydrothermal systems because they are driven primarily by rock-water reactions known as serpentinization, rather than magmatic activity. The serpentinization reactions create high pH fluids that mix with surrounding cold seawater to form the calcium carbonate structures. Serpentinite-hosted ecosystems are of astrobiological interest because they provide a source of fuel and energy for life that does not require sunlight or an active planetary body (i.e. geothermal energy). These systems are thought to be present on inactive planetary bodies such as Jupiter’s moon Europa (1, 2).

The dense microbial biofilms of Lost City chimneys are fueled by the carbon and energy released by serpentinization of the underlying ultramafic rock (3–7). The serpentinization reactions provide high concentrations of hydrogen gas, methane, and other simple organic compounds that serve as food and energy sources for microbes. The more extreme interiors of chimneys are anoxic and continually bathed in the warm serpentinizing fluids. Temperatures of venting fluids can reach >95°C, and pH of the fluids can be as high as 11 (8). Previous studies have shown that these interiors are dominated by a single archaeal phylotype, the Lost City Methanosarcinales (4). In contrast, the chimney exteriors host a more complex microbial community including organisms involved in the oxidation of sulfur and CH_4_ (e.g. *Methylomonas*, *Thiomicroscopira*). These organisms likely thrive in the mixing zones, where they can take advantage of the cooling effect of the seawater and more efficient electron acceptors (e.g. oxygen) but still access the products of serpentinization supplied by venting fluids.

In general, little is known about the metabolic capabilities of Lost City organisms. Our previous research has shown that much of the microbial biomass at Lost City is derived from carbon that originated deep in the Earth’s subsurface (3, 9). In most ecosystems, inorganic carbon (CO_2_) serves as the starting carbon source for primary production. However, the Lost City fluids contain extremely low concentrations of dissolved inorganic carbon (DIC) due to its reduction to hydrocarbons and its rapid precipitation as calcium carbonate at pH above ~9 (10–12). The organic acid formate has been proposed as an alternative primary carbon source; it is present in high concentrations in Lost City fluids (36–158 μM) and has been proposed to form abiotically in serpentinizing fluids (3, 13, 14). In support of this, our previous experiments have shown that the isotopic compositions of carbon (^13^C and ^14^C) in bacterial and archaeal lipids resemble those of formate from the vent fluids (3).

Formate is unable to enter carbon fixation pathways directly and needs to be converted to CO_2_ for autotrophic metabolism. The enzyme formate dehydrogenase catalyzes the reversible oxidation of formate to CO_2_. Formate is too large to passively diffuse through the cell and needs to be brought in through a protein transporter; therefore, any Lost City formate-utilizing species would require both a transporter and a formate dehydrogenase. This study identifies two formate-utilizing species of the Lost City chimneys based on metagenomic evidence, including the presence of formate transporters and formate dehydrogenases. The briefly available CO_2_ leaking from these formate-utilizing species might support the other microbial inhabitants of Lost City chimneys that are unable to use formate, such as the Methanosarcinales.

## RESULTS AND DISCUSSION

### Metagenomic Assembly and Binning

We performed shotgun paired-end sequencing of environmental DNA extracted from a sample of chimney material collected at Marker 5 within the Lost City hydrothermal field. The metagenome consisted of 145,937,844 read pairs (after quality filtering) which were assembled into 730,351 contigs with N50 of 2,518 bp and max length of 250,900 bp. Assembled contigs represent 62.47% of all read pairs in the metagenome. Each contig >1000 bp was assigned taxonomy with PhyloPythiaS+ (15), and the overall taxonomic composition of these contigs is shown in Figure S1. These results are consistent with previous studies that have described the chimney biofilm communities as being dominated by taxa associated with the metabolism of sulfur and methane (e.g. Thiotrichales, Methylococcales, Methanosarcinales) (4–6, 16).

Contigs were binned according to their tetranucleotide frequencies into metagenome-assembled genomes (MAGs), only seven of which initially contained <10% contamination and >18% completeness (***Table S1***) after automated binning. None of these seven MAGs contained strong evidence for formate utilization. Therefore, we manually explored the other MAGs with evidence of formate metabolism.

Formate cannot diffuse through the cell membrane and requires a transporter. In formate-utilizing methanogens, the gene *fdhC* is thought to be necessary for transport of formate into the cell (17). A different formate/nitrite transporter (*focA*) from the same FNT protein family has been described in *Escherichia coli* (18). *E. coli* requires *focA* for the removal of formate produced during mixed-acid fermentations, but the protein is known to be bidirectional and can therefore bring formate, or nitrite, into the cell (19). In order to identify potential formate-using species in our metagenomes, we examined all bins containing *fdhC* or *focA*. We identified five formate transporters in the metagenome, three of which were found on contigs with evidence of nitrite metabolism but no other formate-metabolizing genes. The other two formate transporters were located on contigs with adjacent genes involved in formate metabolism. Therefore, we manually refined these two MAGs, as well as a third representing the Lost City Methanosarcinales phylotype (7, 11).

#### Sulfurovum

The *Sulfurovum* MAG was refined to 95.9% complete and 2.19% contamination by examining the hierarchical clustering of contigs as visualized in anvi’o and by inspecting the taxonomic assignment (by PhyloPythiaS+) of each contig. The fragments mapped to this MAG comprised 0.41% of the total assembly coverage (***Table S2***). Of the 3 MAGs discussed here, the *Sulfurovum* MAG contained the lowest number of protein-encoding genes (2,036), but 90% of these were annotated with a functional prediction. This MAG also had the highest number of complete KEGG modules (53) (***Dataset S1***).

The *Sulfurovum* MAG included a contig with *fdhC* and multiple formate hydrogenlyase (hydrogenase-4, FHL) genes starting 1653 bp away (***Fig. 1***). Interestingly, the formate hydrogenlyase on this contig was homologous to the Hyd-4 form which is unaffected by alkaline pH (20). In *Escherichia coli*, the bidirectional FHL complex links formate oxidation to proton reduction and is operational during mixed-acid fermentations (21). During these fermentations in *E. coli*, formate is formed by the pyruvate formate-lyase enzyme and transported outside of the cell (22). It is unlikely the FHL complex is involved in mixed-acid fermentation by *Sulfurovum* because the MAG did not contain a pyruvate formate-lyase. Therefore, FdhC and the FHL complex most likely bring in formate from the environment and carry out membrane-bound conversion of formate to CO_2_, which can then enter a carbon fixation pathway.

**Figure 1:**
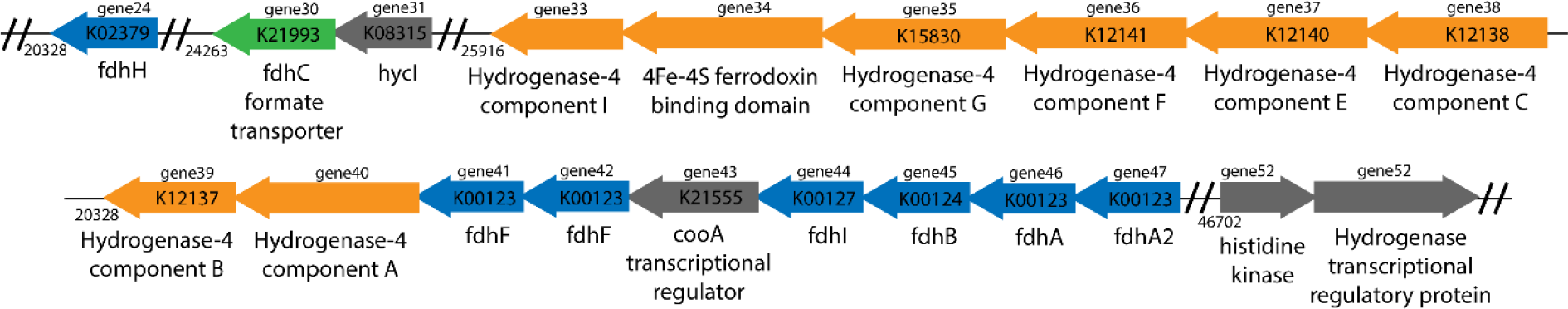
The formate transporter (*fdhC*) contig in the Lost City Sulfurovum MAG. Relevant genes are reported here in the order they are found on the contig. Gene order is indicated above the arrows that indicate forward or reverse direction of transcription. Double lines indicate a break in genes reported. Orange genes are involved in the formate hydrogenlyase (FHL) complex; blue genes (*fdhH*, *fdhF*, *fdhI*, *fdhB*, *fdhA*, *fdhA2*) indicate formate dehydrogenase genes that may interact with the FHL complex; green gene (*fdhC*) is involved in formate transport. All other genes, including regulatory genes, are reported in grey. KEGG id numbers are indicated on arrows where appropriate. *hycI*: hydrogenase 3 maturation protease; *cooA*: CRP/FNR family CO-sensing transcription factor.

Key genes for carbon fixation via the reductive TCA cycle were found in the *Sulfurovum* MAG: pyruvate synthase (PFOR; heterodimer type), ATP citrate lyase, 2-oxoglutarate reductase, and fumarate reductase (***Fig. 2***). In addition, genes for gluconeogenesis were also present, suggesting fixed carbon can be stored as glucose. It is unclear if *Sulfurovum* can run the TCA cycle in the forward direction for heterotrophic growth. Succinate dehydrogenase genes were present in the MAG, but there was no evidence for citrate synthase. *Desulfobacter hydrogenophilus* is known to use ATP citrate lyase (instead of citrate synthase) in both the forward and reverse directions, and this may be possible for *Sulfurovum* as well (23). Alternatively, instead of using a bidirectional TCA cycle to break down glucose reserves, *Sulfurovum* may ferment it into lactate; indeed, lactate dehydrogenase was present in this MAG.

**Figure 2:**
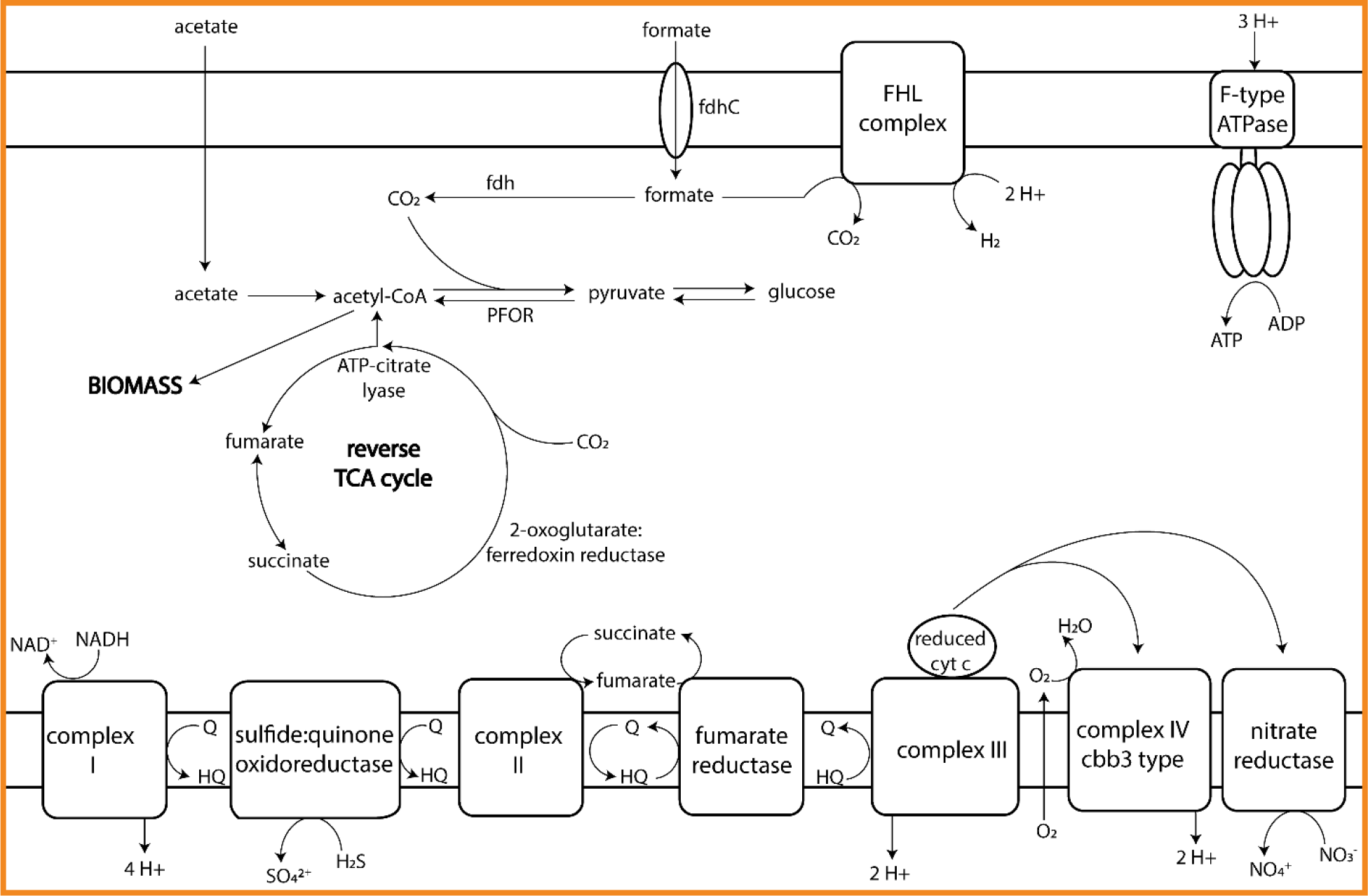
Overview of the Lost City Sulfurovum central carbon metabolism pathways and electron transport chain. Abbreviations: fdh: formate dehydrogenase, fdhC: formate transporter, FHL: formate hydrogenlyase complex, HQ: hydroquinone, Q: quinone, cty c: cytochrome c, PFOR: pyruvate:ferredoxin oxidoreductase

The *Sulfurovum* MAG contained genes for a complex electron transport chain, suggesting a metabolically diverse lifestyle (***Fig. 2***). Complex I (NADH dehydrogenase) likely serves as a versatile entry point for many catabolic reactions. We also found a sulfide:quinone oxidoreductase (SQOR), indicating that *Sulfurovum* can use sulfide as an electron donor when present in the environment. The electron transport chain found in this MAG contained three complexes that can serve as terminal electron acceptors: fumarate reductase, complex III and cytochrome c oxidoreductase (complex IV), and nitrate reductase.

This MAG also contained a number of cofactor ABC transporters including those for tungstate (required for formate dehydrogenase activity), molybdate, iron, thiamin, and zinc. We also found evidence of transporters for macronutrients such as L-amino acids, branched-chain amino acids, phospholipids and phosphate.

In addition to these nutrient-acquiring transporters, the MAG contained a transporter responsible for excreting capsular polysaccharides. After intracellular construction, these molecules are exported to form a capsule around the cell which is involved in both biofilm formation and environmental stress protection (24). We also found a lipopolysaccharide export system indicating that the *Sulfurovum* species at Lost City builds an outer membrane like other characterized *Sulfurovum* species (25).

*Sulfurovum lithotrophicum*, the type species for the genus, was first isolated from hydrothermal sediments off the coast of Okinawa, Japan (26). The genome for this species closely resembles our Lost City *Sulfurovum* MAG. It contains a rTCA cycle, sulfide:quinone oxidoreductase (SQOR), and can use O_2_ or NO_3_^-^ as an electron acceptor (27). However, unlike our *Sulfurovum* MAG, this and most characterized *Sulfurovum* species are also able to oxidize sulfur (S^0^) or thiosulfate through the sulfur-oxidation (SOX) system (28–30). Both *Sulfurovum aggregans* and *Sulfurovum lithotrophicum* have been shown to use hydrogen, but not formate, as an electron donor (29, 31).

The lack of a SOX system and the presence of formate-metabolizing genes in the Lost City *Sulfurovum* MAG are novel for the genus. The ability to scavenge amino acids, form biofilms, and retain genetic flexibility for multiple electron acceptors supports this organism having a mixotrophic lifestyle capable of adapting to a fluctuating environment with varying ratios of seawater and hydrothermal fluids. These results suggest that the ideal location for *Sulfurovum* is in a transition zone between the interior and exterior of the chimneys.

### Chloroflexi

After refining, the Chloroflexi MAG is estimated to be 70% complete with 4.73% contamination. The mapped fragments make up 0.62% of the assembly (***Table S2***). Of the three MAGs discussed here, this MAG contained the highest number of protein-encoding genes (3,936), but only 84% of these were annotated with functional predictions. This bin also had the highest number of incomplete KEGG modules (39) (***Dataset 1***).

The Chloroflexi MAG contained a *focA* formate transporter adjacent to a formate dehydrogenase alpha subunit (*fdhA*) gene and three genes encoding the catalytic subunit of NAD(H) dehydrogenase (*nuoG*, *hoxF*, *hoxE*) (***Fig. 3***). The beta subunit (*fdhB*) was located together with *fdhA* on a different contig. In addition to formate-utilizing genes, the Chloroflexi MAG contained a carbon fixation pathway. A nearly complete reductive pentose phosphate cycle was present (***Fig. 4***). The MAG also included a carboxysome-specific carbonic anhydrase, suggesting that this organism uses a carboxysome to concentrate CO_2_ around the ribulose-1,5-bisphosphate carboxylase/oxygenase (RuBisCO) (32). Carboxysomes are used by many organisms when the concentration of CO_2_ outside the cell is lower than the K_m_ of RuBisCO (33). If the Chloroflexi species couples the carboxysome shell with the conversion of formate to CO_2_, this could be an effective adaptation to the lack of CO_2_ in the chimney environment.

**Figure 3:**
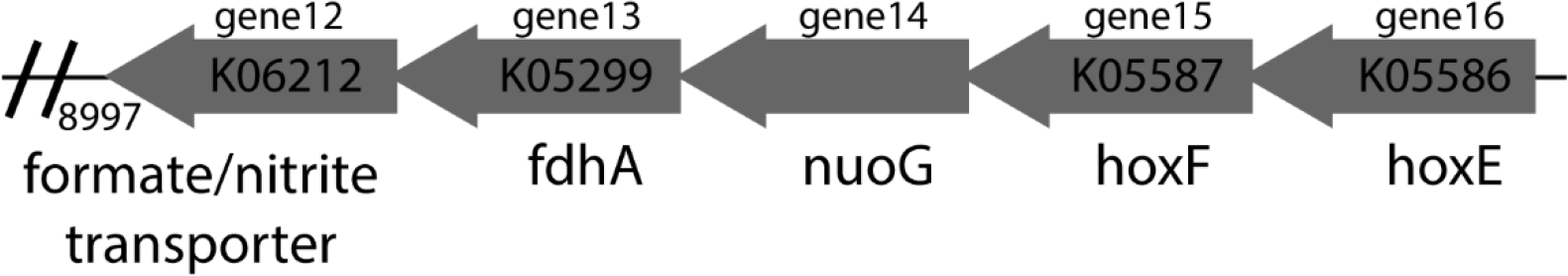
The formate transporter (*focA*) contig found in the Lost City Chloroflexi MAG. Note the adjacent gene encoding the alpha subunit of formate dehydrogenase (*fdhA*). Double lines indicate a break in genes reported. KEGG id numbers are indicated on arrows where appropriate. *nuoG*: NADH-quinone oxidoreductase subunit G; *hoxF* & *hoxE*: bidirectional [NiFe] hydrogenase diaphorase subunit.

**Figure 4:**
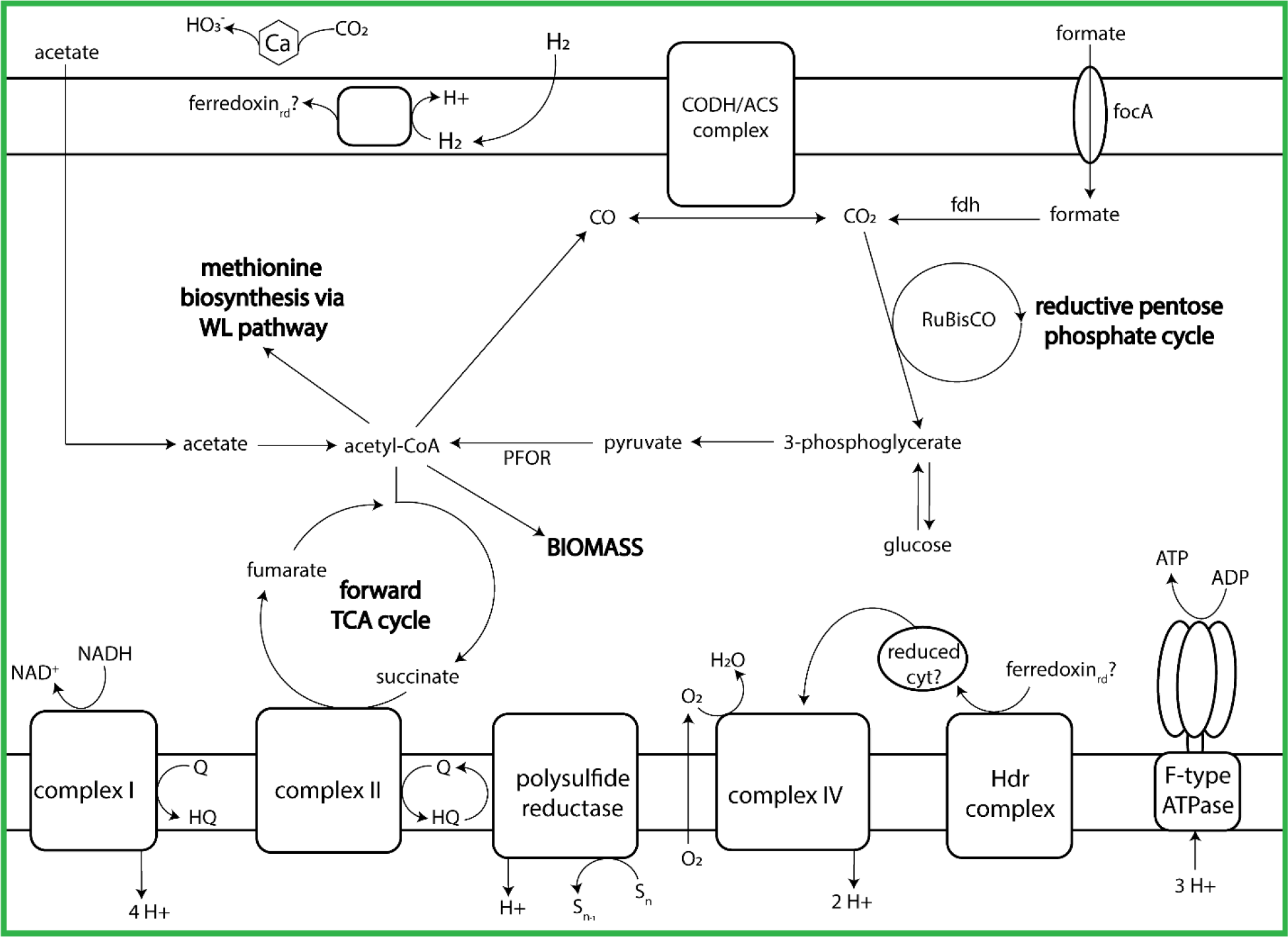
Overview of the Lost City Chloroflexi central carbon metabolism pathways and electron transport chain. Abbreviations: RuBisCO: ribulose-1,5-bisphosphate carboxylase/oxygenase, Hdr: heterodisulfide reductase, fdh: formate dehydrogenase, CODH/ACS: carbon monoxide dehydrogenase/acetyl-CoA synthase, focA: formate/nitrite transporter, WL pathway: Wood-Ljungdahl pathway, Ca: carbonic anhydrase, cty: cytochrome, PFOR: pyruvate:ferredoxin oxidoreductase, HQ: hydroquinone, Q: quinone, rd: reduced

The one enzyme missing from the reductive pentose phosphate cycle in this MAG is glyceraldehyde-3-phosphate dehydrogenase (GAPDH), which is also involved in glycolysis. Interestingly, thermophilic organisms use a distinct form of GAPDH to cope with the heat instability of glyceraldehyde-3-phosphate (23, 34), so it is possible that the Chloroflexi MAG contains an as yet unidentified variant of GAPDH. Another adaptation of thermophilic growth is in the structure of fructose-bisphosphate aldolase, an enzyme involved in gluconeogenesis (34). This gene is also included in the Chloroflexi MAG and is most closely related to that of the thermophile *Caldilinea aerophila*.

The Chloroflexi MAG also contained an incomplete Wood-Ljungdahl pathway. Typically, the presence of this pathway in a bacterial genome indicates that it is involved in carbon fixation during acetogenesis, but the lack of two key enzymes in this MAG casts doubt on that scenario. One of these missing enzymes (methylene-tetrahydrofolate reductase: MTHFR) is essential for acetogenesis. The second missing enzyme is acetate kinase, which is involved in the last step of acetate formation. A similar partial Wood-Ljungdahl pathway has been described in the dehalogenating Chloroflexi *Dehalococcoides mccartyi* (35). Although *D. mccartyi* is missing MTHFR, the species is capable of *de novo* methionine biosynthesis through the partial Wood-Ljungdahl pathway via the cleavage of acetyl-CoA. As the Lost City Chloroflexi MAG contained no other pathways for methionine biosynthesis, this organism, like *D. mccartyi,* may use the incomplete Wood-Ljungdahl pathway for methionine biosynthesis rather than for carbon fixation.

The Chloroflexi MAG had both glycolysis/gluconeogenesis pathways and a forward TCA cycle. This suggests the organism could store carbon fixed through the reductive pentose phosphate pathways as glucose reserves and grow heterotrophically when carbon is limited in the environment. Additional evidence for a flexible mixotrophic lifestyle for this MAG included three carbohydrate transporters (multiple sugar, ribose, and D-xylose transport systems), suggesting that this Chloroflexi species is capable of metabolizing additional complex carbon sources.

The membrane-bound complexes encoded by the Chloroflexi MAG are consistent with an anaerobic, mixotrophic lifestyle (***Fig. 4***). Energy conservation appears to be mediated by a complete NADH:quinone oxidoreductase (complex 1) (*nuoABCDEFGHIJKLMN*), a succinate dehydrogenase (complex II), a polysulfide reductase, a cytochrome c oxidase (complex IV), a heterodisulfide reductase, and an F-type ATPase typical of bacteria. Electrons could be donated by formate or carbohydrates, but the terminal electron acceptor is unclear.

The MAG had a cytochrome c oxidase (complex IV), which would indicate oxygen as the terminal electron acceptor, but the lack of a cytochrome oxidoreductase such as cytochrome bc or b6f complex (complex III) is perplexing. The genome of anaerobic *Sulfurimonas gotlandica* str. GD1 contains a cytochrome c oxidase, but the enzyme’s suspected function is to occasionally remove inhibitory oxygen rather than to serve as a terminal electron acceptor (36). Furthermore, the Chloroflexi genus *Anaerolineae* is described as obligately anaerobic, yet many species contain genes for aerobic respiration in their genomes (37). Due to the presence of numerous oxygen-sensitive enzymes (aldehyde ferredoxin oxidoreductase, CODH/ACS complex, anaerobic forms of glycerol-3-phosphate dehydrogenase and sulfatase maturase) in the Chloroflexi MAG, it is likely that the cytochrome c oxidase’s role is to remove oxygen rather than to serve in the last step of an aerobic respiratory chain.

However, the lack of a complex III raises the question of where the reduced cytochrome would be transferred from. Heterodisulfide reductase (Hdr) has been proposed to be involved in electron transfer to cytochromes in other species (38, 39). This complex could work in tandem with the cytochrome c oxidase to remove intracellular oxygen. If oxygen is not the terminal electron acceptor, then the presence of polysulfide reductase (*Psr*) indicates polysulfide compounds as the most likely terminal electron acceptors. The Chloroflexi polysulfide reductase sequence has 40% identities with the *Psr* sequence from *Thermus thermophilus*, which has been shown to use polysulfide as its terminal electron acceptor (40).

The Chloroflexi MAG contained transporters for tungstate (required for formate dehydrogenase activity), iron, and thiamin. As with the *Sulfurovum*, the Chloroflexi is able to scavenge some macronutrients: L-amino acids, branched-chain amino acids, phospholipids and phosphate. Because the MAG is estimated to be only 70% complete, it is likely that it contains additional transporters not identified in this study.

Due to the presence of oxygen-sensitive enzymes, the Lost City Chloroflexi species is expected to be an anaerobe which lives in or very close to the interior of the chimneys. Carbon from formate can be converted into biomass, which could lead to CO_2_ leaking from the cell and fueling autotrophic species like the Lost City Methanosarcinales.

### Methanosarcinales

The refined Methanosarcinales MAG was estimated to be 84.87% complete with 5.26% contamination. The mapped fragments made up 4.41% of the entire assembly (***Table S2***). The MAG contained 2,324 protein-encoding genes, only 77% of which could be assigned a predicted function. The Methanosarcinales MAG had 33 complete and 28 incomplete KEGG modules (***Dataset S1***).

We identified this MAG as the previously described Lost City Methanosarcinales phylotype due to the taxonomic assignment of Methanosarcinales for all contigs and the presence of nitrogenase reductase (*nifH*, nitrogen fixation) and methyl coenzyme M reductase (*mcrA*, methanogenesis) sequences that matched those of previously sequenced genes (7, 11) (***Table S3, S4***). In agreement with our previous analysis with a smaller dataset, we found no evidence that the Methanosarcinales is able to utilize formate as a carbon source (3). The MAG did contain a genomic inventory that would allow the Methanosarcinales to utilize CO_2_, acetate, and methanol for methanogenesis (***Fig. 5***). Transporters for tungstate and molybdate were also identified; these metals are required cofactors for many of the enzymes in the methanogenic pathway.

**Figure 5:**
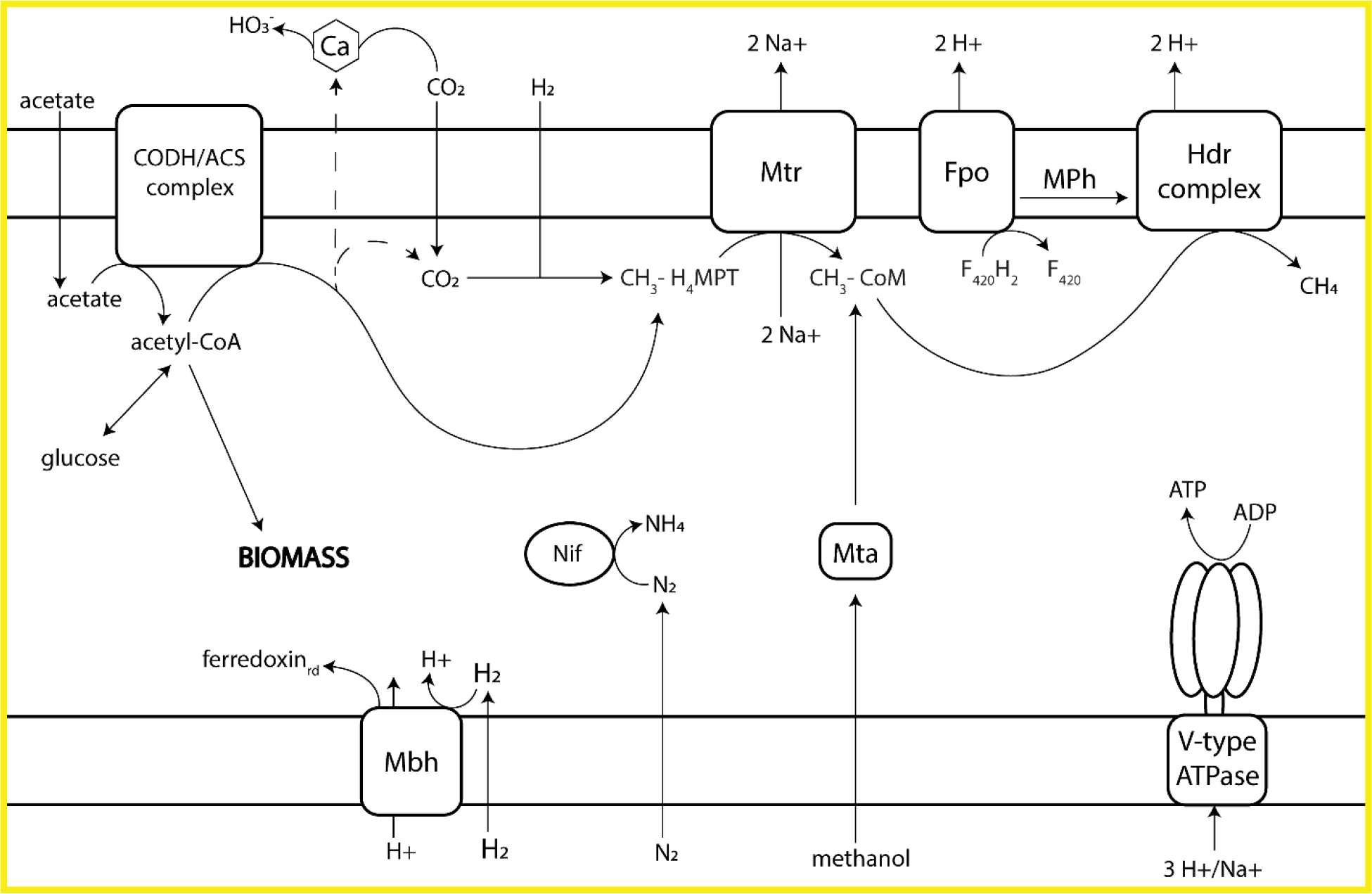
Overview of the Lost City Methanosarcinales (LCMS) methanogenic pathway and membrane-bound complexes involved in energy conservation. Abbreviations: CODH/ACS: carbon monoxide dehydrogenase/acetyl-CoA synthase, Mbh: membrane-bound hydrogenase, Nif: nitrogenase, Mtr: methyl-H4MPT:coenzyme M methyltransferase, Mta: methanol:5-hydroxybenzimidazolylcobamide methyltransferase, Fpo: F_420_H_2_ dehydrogenase, MPh: methanophenazine, Hdr: heterodisulfide reductase, rd: reduced

This Methanosarcinales MAG provides new information on the phylogenetic status of this species. Previous studies classified this species within order Methanosarcinales (4), but this organism has never been cultured, despite many attempts, and a more specific phylogenetic classification has never been attempted. Its closest relatives have been previously reported to include members of the *Methanosarcinaceae* and *Methanosaetaceae* families (4, 6). These two families include the only species known to be capable of methane production from acetate (41). Each group of methanogens has distinct mechanisms for acetate activation and energy conservation. Acetate activation in *Methanosarcina* species proceeds via two enzymes: acetate kinase and phosphotransacetylase. *Methanosaeta* species, by contrast, use acetyl-CoA synthetase (ACS) for acetate activation. The LCMS MAG included both ACS and acetate kinase (but not phosphotransacetylase, perhaps due to the incompleteness of the MAG), suggesting that it may be able to use both systems or a hybrid system.

The energy conservation strategy of the Lost City Methanosarcinales appears to be more similar to that of *Methanosaeta* than *Methanosarcina*. As in *Methanosaeta*, the MAG contained no Ech hydrogenase, no Rnf complex, and no methanophenazine-reducing hydrogenase (Vho). Instead, the only energy-conserving complex external to the methanogenesis pathway is F_420_H_2_ dehydrogenase, which is also employed by *Methanosaeta* species (41).

Phylogenetic analyses of methanogenesis genes from this MAG suggest that it is distinct from both *Methanosaeta* and *Methanosarcina*, perhaps forming a novel family within order Methanosarcinales (***Figure S2***). In two of the gene trees (*ftr* and *mer*), the Lost City gene was monophyletic with methylotrophic methanogens such as *Methanococoides burtonii*. A third phylogeny (*mtd*) grouped the Lost City gene with hydrogenotrophic and formate-utilizing Methanosarcinales, such as *Methanothermococcus okinawensis*. In a fourth phylogeny (mch), the Lost City gene was distinct from all other known species.

No genomes of *Methanosaeta* contain membrane-bound hydrogenases that would allow H_2_ to serve as the electron donor (42–44). However, the Lost City MAG contained five genes annotated as a ferredoxin-dependent membrane bound hydrogenase (Mbh). This complex is known to translocate protons with the formation or cleavage of hydrogen gas, similar to the Ech hydrogenase found in many hydrogenotrophic methanogens (45). Therefore, the Lost City Methansarcinales may be able to use this enzyme for hydrogenotrophic methanogenesis.

For hydrogenotrophic growth, the Lost City Methanosarcinales would be reliant on CO_2_ leaking from a formate-utilizing species (such as *Sulfurovum* or Chloroflexi), but the relationship could be mutualistic. The Methanosarcinales MAG contained nitrogenase genes, responsible for nitrogen fixation. Previous work found low δ^15^N values of Lost City chimneys, indicative of biological nitrogen fixation (46). The concentrations of biologically available nitrogen are relatively low (<6 uM) in Lost City fluids, but concentrations of N_2_ resemble that of seawater (46). Therefore, the densely populated biofilm communities of Lost City chimneys must be reliant on nitrogen fixation, probably carried out at least in part by the species represented by the Methanosarcinales MAG.

## CONCLUSION

The biofilms growing on Lost City chimneys are unique ecosystems where microbes must face the challenges of multiple extremes, including pH >10 and temperatures up to at least 95°C. Our previous work demonstrated that microbial communities inhabiting the chimneys are fueled by carbon venting from Earth’s mantle (3). This work identifies Chloroflexi and *Sulfurovum* species as the formate-utilizing organisms that may be required to make mantle-derived carbon available to the rest of the chimney ecosystem.

The Lost City biofilms that inhabit the anoxic interiors of the chimneys have been described as containing a single-species, the Lost City Methanosarcinales (3–7, 11). The single Methanosarcinales MAG reported here represents 4.41% of the chimney metagenome, >5 times more abundant than the other MAGs we have been able to recover so far. This species has been previously shown to dominate the chimney microbial community (4, 5, 7), yet it appears to be unable to use one of the most abundant carbon sources, formate. The Lost City Chloroflexi MAG, by contrast, contains the required genes for using formate and may provide CO_2_ to Methanosarcinales and other members of the biofilm community. The Chloroflexi MAG is seven times less abundant than the Methanosarcinales MAG, so it appears to serve its carbon-cycling role as a low-abundance but foundational member of the community. Alternatively, the Chloroflexi species may be more abundant in the rocky subsurface of the Atlantis Massif, where it can convert mantle-derived formate into more biologically accessible CO_2_ for surface communities. Future research should investigate how microbial activity in subsurface environments underlying the Lost City might influence the food and energy available to the biofilm communities of the chimneys.

## METHODS

### Sample Collection

Sample H08_080105_Bio5slurpB1 from Marker 5 was collected in 2005 during a National Oceanic and Atmospheric Administration (NOAA) Ocean Explorer cruise with the ROV Hercules aboard the R/V Ronald H. Brown. The sample was immediately placed in a sterile Whirl-Pak® sample bag upon arrival on deck and stored at −80 °C until analysis. DNA was extracted from the samples according to a previously published protocol (5). A previous metagenomic analysis of this sample has been published, (3) but the results presented here are from a different DNA extraction and a much deeper metagenomic sequencing effort.

Environmental DNA was extracted with a protocol modified from (4, 6, 47). The chimney sample was crushed and homogenized with a sterile mortar and pestle, and 0.25-g subsamples were placed in a DNA extraction buffer containing 0.1 M Tris, 0.1 M Na-EDTA, 0.1 M KH_2_PO_4_, 1.5M NaCl, 0.8 M guanidium HCl, and 0.5% Triton X-100. For lysis, samples were subjected to one freeze-thaw cycle, incubation at 65°C for 30 min, and beating with 0.1-mm glass beads in a Mini-Beadbeater-16 (Biospec Products). Purification was performed via extraction with phenol-chloroform-isoamyl alcohol, precipitation in 3M sodium acetate and ethanol, washing in Amicon 30K Ultra centrifugal filters, and final cleanup with 2× SPRI beads (48). DNA quantification was performed with a Qubit fluorometer (Thermo Fisher).

### Metagenome Sequencing

A metagenome library was constructed using the NEBNext Ultra DNA Library Prep Kit for Illumina according to the manufacturer’s instructions. Quality control and sequencing of the metagenomic libraries was conducted at the University of Utah High-Throughput Genomics Core Facility. Libraries were evaluated for quality on a Bioanalyzer DNA 1000 chip (Agilent Technologies), and then paired-end sequencing (2 × 125 bp) was performed on an Illumina HiSeq2500 platform with HiSeq v4 chemistry. The library was multiplexed with one other library (from a second Lost City chimney sample; results from which are not reported here) on one Illumina lane, yielding 180 million read pairs (45 billion bases). Demultiplexing and conversion of the raw sequencing base-call data was performed through the CASAVA v1.8 pipeline.

### Metagenomic Processing

Raw sequence data was processed by the Brazelton lab to trim adapter sequences with BBDuk (part of the BBTools suite (49)), to remove artificial replicates, and to trim reads based on quality. Removal of replicates and quality trimming were performed with our seq-qc package (https://github.com/Brazelton-Lab/seq-qc). Paired-end reads were assembled with MegaHit using kmers of 27-141. Prodigal was run in the anonymous gene prediction mode to identify ORFs. Functional annotation was performed using Diamond blastp and using both the KEGG-T10000 and KEGG-prokaryotes databases with an e-value of 1e-6. Annotations were selected by highest quality alignment as determined by the bit score. Binning was performed with ABAWACA (https://github.com/CK7/abawaca). Contig taxonomy was assigned with PhyloPythiaS+ (15). Curation of bins was performed in anvi’o, using the default visualization of bins informed by tetranucleotide frequency as well as manual inspection of PhyloPythiaS+ taxonomic assignment. CheckM was used to assess bin quality (50). Completion of KEGG modules and pathways was determined using the KEGG Mapper online tool (https://www.genome.jp/kegg/mapper.html). Coverage was determined through read mapping with bowtie2 and bedtools genomecov (51, 52). Reference proteins for phylogenetic trees were downloaded from NCBI GenBank in September 2019 and can be found in Table S5. The multiple sequence alignments were conducted with MUSCLE (53), and the phylogenies were built using RaxML and the “-f a” option with 100 bootstrap replicates.

## Data Availability

All unassembled sequence data related to this study are available at the NCBI Sequence Read Archive (BioSample: SAMN13035994) and MAG assemblies have been submitted to NCBI GenBank (accession numbers pending). All NCBI data is under BioProject PRJNA577730. All SRA metadata, full supplementary material, and protocols are archived at https://doi.org/10.5281/zenodo.3522424. All custom software and scripts are available at https://github.com/Brazelton-Lab.

## Acknowledgements

The authors would like to thank Emily Dart for DNA extractions and metagenomic library preparation, Alex Hyer and Christopher Thornton for computational assistance, and Dr. David Blair for assistance with metabolic pathways. Funding was provided by the National Science Foundation (NSF projects 1536702 and 1536406), the Center for Dark Energy Biosphere Investigations, and the NASA Astrobiology Institute Rock-Powered Life team.

## Supplemental Files

**Table S1:** Other MAGs greater than 18% complete and under 10% contamination as determined by CheckM. PhyloPhythiaS+ taxonomic consensus is reported if one third or more of the contigs in the MAG were assigned the same taxonomic grouping.

**Table S2:** Assembly quality statistics for each quality MAG.

**Table S3:** BLAST results for previously sequenced *nifH* genes.

**Table S4:** BLAST results for previously sequenced *mcrA* genes.

**Table S5:** Accession numbers for reference proteins used in Figure S2.

**Dataset S1:** DatasetS1.xlsx - Complete and incomplete KEGG modules for the Methanosarcinales, Sulfurovum, Chloroflexi MAGs.

**Figure S1:** Coverage of taxonomic assignments of contigs in the assembly. Taxonomy is reported at the level of order. Exceptions are *Sulfurovum* and contigs with unassigned order, where lowest assignment is reported.

**Figure S2**: Phylogenetic trees of Lost City Methanosarcinales (LCMS) MAG genes in methanogenic pathways. **a.** formylmethanofuran-tetrahydromethanopterin formyltransferase (*ftr*), **b.** N^5^N^10^-methenyl-tetrahydromethanopterin cyclohydrolase (*mch*), **c.** F_420_-dependent N^5^N^10^-methylene-tetrahydromethanopterin dehydrogenase (*mtd*), **d.** and F_420_-dependent N^5^N^10^-methylene- tetrahydromethanopterin reductase (*mer*). Accession numbers for reference proteins can be found in Table S5.

